# Photo-excited Toluidine Blue disaggregates the Repeat Tau and modulates cytoskeletal structure in neuronal cells

**DOI:** 10.1101/2020.03.06.980276

**Authors:** Tushar Dubey, Nalini Vijay Gorantla, Subashchandrabose Chinnathambi

## Abstract

Alzheimer’s disease is a progressive neurological disorder characterized by the intracellular accumulation of Tau protein aggregates. Inhibition of protein aggregation by photo-excited dyes is emerging as novel strategy for the treatment of certain diseases. Toluidine Blue is a basic phenothiazine dye having potency of photo-excitation by irradiation with red light at 630±20 nm. In present work, we studied the effect of Toluidine Blue and photo-excited TB on aggregation of repeat Tau in-*vitro* using Thioflavin S fluorescence assay, SDS-PAGE and electron microscopy. Results show that TB efficiently inhabited the formation of higher order aggregates. Moreover, the photo-excited TB led to disaggregation of the mature repeat Tau fibrils. Further, studies on the effect of Toluidine blue on cell viability and cytoskeleton network of Neuro2acells show that TB was not toxic to neuronal cells at lower concentrations but at high concentrations (> 5 μM) both TB and photo-excited TB induced significant toxicity. Immunofluorescence studies on the cytoskeleton of Neuro2a cells show that Toluidine Blue and photo-excited Toluidine Blue treatment at non-toxic concentration of 0.5 μM stimulated formation of actin rich lamellipodia and filopodia structures. Tubulin networks were also differentially modulated after the treatment of Toluidine Blue and photo-excited Toluidine Blue. End Binding protein 1 (EB1) levels were observed to increase after Toluidine Blue and photo-excited Toluidine Blue treatment indicating the accelerated microtubule polymerization. The overall study suggested that Toluidine Blue inhibited the aggregation of soluble Tau and photo-excited Toluidine Blue disaggregated the pre-formed Tau filaments.

## Introduction

Alzheimer’s disease is characterized by accumulation of intracellular Tau in the form of neurofibrillary tangles (NFTs)(Mandelkow & Mandelkow 2012; Gorantla *et al.* 2017). Apart from AD, the aggregated Tau has been reported to be involved in several other diseases including Pick’s disease, corticobasal degeneration, progressive supranuclear palsy, post encephalitic parkinsonism *etc.*, grouped as Tauopathies(Iqbal *et al.* 2005; Sonawane & Chinnathambi 2018). Tau is a natively unfolded protein present in cytoskeleton having physiological role in stabilization of the microtubules and cytoskeleton structure(Mitchison & Cramer 1996). In neurodegenerative diseases, Tau aggregation has been reported to be closely associated with cytoskeleton abnormalities in neuronal cells involving the actin and tubulin deformities(Bamburg & Bloom 2009; Cairns *et al.* 2004). Actin and Tubulin are the abundant cytoskeleton proteinsTubulin is known to be involved in formation of microtubules whereas, actin majorly assist the cells in substratum adhesion, synapse formation and cell motility by the formation of structures as lamellipodia, filopodia, podosomes *etc*(Hadfield *et al.* 2003; Small *et al.* 2002). Similarly the End Binding protein 1 (EB1) is a cytoskeletal associated protein which has been reported to be located at growing end of microtubules(Stepanova *et al.* 2003). The hexapeptide VQIINK and VQIVYK present in 2^nd^ and 3^rd^ repeat respectively, are reported to to be involved in Tau aggregation(Ballatore *et al.* 2007; Balmik & Chinnathambi 2018). To study the aggregation inhibition propensity of certain molecules, the repeat region of Tau has been used as a target protein(Gorantla *et al.* 2019b; Khlistunova *et al.* 2006). Tau is considered as phospho-protein, which requires phosphorylation to perform its activity, but under pathological conditions hyperphosphorylation Tau by various kinases e.g. *GSK*3β and CDK5 leads to generation of Tauopathy(Das *et al.* 2020; Sonawane *et al.* 2019b). Several compounds of natural and synthetic origin have been studied against Tau aggregation(Gorantla *et al.* 2019a; Pickhardt *et al.* 2005). These compounds can be categorised in two groups based on their action firstly disaggregation of the aggregated Tau and second inhibition of the aggregation of Tau e.g as reported for anthraquinones and EGCG respectively(Sonawane *et al.* 2019b). Several dyes such as methylene blue, TBO and rose bangal have been reported to modulate β-Amyloid Peptides aggregation (Lee *et al.* 2017; Lee *et al.* 2015; Sonawane *et al.* 2019b). Phthalocyanine dye has been reported to be effective against pathological prion protein (PrPC)(Kostelanska *et al.* 2019). Moreover photo-active chlorin 6 dye was also found to reduce the aggregation by modulating the histidine residues(Leshem *et al.* 2019). Methylene blue derived Leuco-methylthioninium Bis (Hydromehanesulphonate) (LMTM) entered phase-3 clinical trial for treatment of mild Alzheimer’s disease(Wilcock *et al.* 2018), however LMTM in further studies was found to be ineffective(Sun *et al.* 2018). Toluidine Blue (TB) is a basic phenothiazine dye having structural similarity with methylene blue. In present study, we evaluated the potency of TB against repeat Tau aggregation. Furthermore, the effect of TB in presence of irradiation was also studied in disaggregating the matured fibrillary aggregates of repeat Tau. In present study the potency of TB and PE-TB against Tauopathy was studied in various aspects.

## Materials and Methods

### Materials

MES (M3671), BSA (82516), BES (14853), BCA (B9643), CuSO_4_ (C2284) ThS (T1892), Toluidine Blue (T3260), MTT (M2128) were purchased from Sigma. IPTG (420322) and DTT (3870) were purchased from Calbiochem. Other chemicals such as Ampicillin (2007081), NaCl (194848), KCl (194844), Na_2_HPO_4_ (191437), KH_2_PO_4_ (19142), EGTA (194823), MgCl_2_ (191421), PMSF (195381), Ammonium acetate (191404) and Heparin (904108), DMSO were from purchased from MP Biomedicals and protease inhibitor cocktail was from Roche. Copper coated carbon grids were purchased from Ted Pella (01814F, carbon type-B, 400 mesh, Cu) DMEM advanced F12 media (12634010), Fetal Bovine serum (16000044), Pensterp cocktail (04693159001), Anti-anti (15240062) were purchased from Gibco.

### Purification of recombinant Repeat Tau

The recombinant Tau expressed in *E.coli* BL 21* cells was purified by the method suggested in earlier literature (Gorantla *et al.* 2019a). Briefly, the *E.coli* culture was grown in Luria-Bertani broth at 37°C, at 180 rpm in orbital shaker (INFORS HT) till culture obtained OD_600_ of 0.5. In log phase the culture was induced with 0.5 mM isopropyl β-D-1-thiogalactopyranoside (IPTG) and incubated for further 4 hours at 37 °C in orbital shaker at 180 rpm. Further, 4 hours of post-induction incubation time was followed according to standardized protocol (Sonawane *et al.* 2019a). The culture was pelleted down by centrifuging at 4000 rpm for 10 minutes in Avanti JXN 26. The pellets were further suspended in lysis buffer (50 mM MES, 1 mM EGTA, 2 mM MgCl_2_, 5 mM DTT, 1 mM PMSF and 50 mM NaCl), the lysis was carried out under high pressure. The culture suspension was lysed at 15000 psi of pressure in homogenizer (Constant cell disruption system) with continuous lysis cycle. After 2 cycles of lysis the lysate was collected and stores on ice for reducing the protein degradation. The subsequence step of lysis is the lysate heating, the lysate was heated at 90°C for 20 min in water bath (Benchmark) without any agitation. After heating the lysate was allowed to cool and then centrifuged at 164,700 × g (Optima XPM ultracentrifuge, Beckman Coulter) for 45 min at 4°C. The supernatant was collected and dialyzed against buffer A. Tau is positively charged protein, hence the purification was done by cation-exchange chromatography as described in previous studies (Gorantla *et al.* 2017). The Repeat Tau was eluted by giving a gradient of NaCl, the fraction under the elution peak were collected and concentrated. The desired molecular weight of our protein was 13.4 kDa, hence the quality of protein was observed on 17 % sodium dodecyl sulfate–polyacrylamide gel (SDS-PAGE). The concentration of protein was estimated by bicinchoninic acid assay (BCA assay) (Barghorn *et al.* 2005). Briefly, bovine serum albumin (BSA) was diluted for making the standard graph. Tau protein was diluted in ratios of (1:100, 1:200, and 1:400). BCA reagent was prepared freshly by mixing bicinchoninic acid and CuSO_4_ in the ratio of 4:1. After addition of BCA to protein, the mixture was incubated at 37°C for 60 minutes in dark. The absorbance was measured at 562 nm in Tecan M 200 multimode plate reader and concertation was estimated referring standard graph.

### Preparation of repeat Tau-fibrils

The soluble repeat Tau was induced to form aggregates by incubating with heparin as suggested in published literature(Sonawane *et al.* 2019a; Sonawane *et al.* 2019b). 100 μM of soluble repeat Tau was incubated with 25 μM heparin. The assembly was carried out in 20 mM BES buffer supplemented with 1 mM DTT, 25 mM NaCl, and protease inhibitor cocktail. The assembly mixture was kept at 37°C for 3 days and the aggregates were confirmed by SDS-PAGE and electron microscopy. The heterogeneous Tau aggregates were observed on SDS-PAGE as bands of higher molecular weights(Gorantla *et al.* 2018). The higher order bands (25-150 kDa) were observed for the samples of aggregates. Moreover the electron microscopy images suggested the presence of long tangled Tau filaments, which supported the presence of aggregates in our sample.

### ThS fluorescence assay

Thioflavin S (ThS) assay was used to monitor the assembly of Tau fibrils. The procedure was followed according to earlier published work (Sonawane *et al.* 2019b). The Tau protein was measured at a concentration of 2 μM incubated with 8 μM ThS dissolved in ammonium acetate. The mixture was incubated at room temperature in dark for 10 min and the fluorescence was measured at excitation wavelength of 480 nm and emission was measured at 520 nm in Tecan M 200 multimode plate reader.

### Electron microscopy

The morphological analysis of Tau fibrils was done by scanning under transmission electron microscopy. Tau was incubated at a concentration of 2 μM on 400 mesh copper coated carbon grid for 90 seconds. After 2 subsequent washes with milliQ the grid was incubated with 2% uranyl acetate for 120 seconds. The grids were blot dried and kept at room temperature. The scanning was done on Tecnai 20 electron microscope.

### Photo-irradiation of TB

Toluidine Blue has an absorption maxima of 630 nm, thus the TB was photo-excited by red light. For photo-excitation a dark chamber was designed, which comprises of commercially procured red LED light source (3.5 watt). The setup was equipped with thermometer to measure heat changes during the irradiation. The repeat Tau aggregates were mixed with various concentrations of TB (2-40 μM). The mixture was transferred to 96 well black well plate. The plate was exposed to red light in dark chamber for the photo-excitation of TB. After 180 minutes of irradiation, the samples were taken out from the plates and analysed by various biochemical and biophysical assays including SDS-PAGE, ThS fluorescence assay and electron microscopy. The irradiation dose or irradiance was calculated by formula

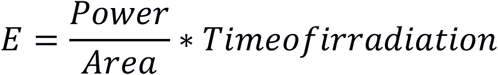

Where E is irradiance, time was calculated in terms of seconds.

### Cell biology studies

Mouse neuroblastoma cells Neuro2a (ATCC: CCL-131), were cultures in advanced DMEM F12 media. 10^4^ cells/well were seeded in 96 well plate for the viability assay. The cells were treated with various concentrations of TB viz, 1, 2.5, 5, 10, 20, 40, 80 and 120 μM for 24 hours. After the incubation 10 μl of 5 mg/ml MTT solution was added in each well. The plate was further incubated at 37°C for 4 hours. The formazan crystals were dissolved in 100% DMSO and the absorbance was measured at 570 nm.

### Immunofluorescence analysis

Neuro2a cells (50,000/well) were seeded on glass coverslip and allowed to incubate for 24 hours at 37°C. The cells were treated with various concentrations of TB (0.5 and 50 μM) and irradiated for 10 minutes with red light. After 24 hours of incubation with PE-TB, the cells were processed for immunostaining. The treated and untreated cells were fixed with absolute methanol for 20 minutes at −20°C. 0.2% Triton X-100 was used for cell permeabilization. For avoiding the non-specific binding of antibody, the cells were incubated with 5% horse serum for 1 hour. The cells were incubated with anti-tubulin (Thermo PA1-41331; dilution 1:250) and K9JA (Dako A0024; dilution 1:500) antibody. After overnight incubation the cells were incubated secondary antibody tagged with Alexa Fluor 488 (Thermo, A11034) (anti-rabbit; dilution 1:1000) and Alexa Fluor 555 (Thermo, A32727) (Anti-mouse; dilution 1:500). The nucleus was stain with DAPI. After the incubation coverslips were mounted using mounting media (70% glycerol) and were sealed on a glass slide. These slides were allowed to air dry at room temperature before the scanning. These samples were scanned on Zeiss Axio observer inverted microscope using 63X magnification in oil emersion and at 40% light intensity.

### Disaggregation of PE-TB

TB treated repeat Tau aggregates were irradiated with red light for 180 minutes. The PE-TB treated aggregates and untreated samples were incubated with 8 μM of ThS dye (diluted in 50 mM ammonium acetates) for 10 minutes in dark conditions. The samples were transferred in 384 black well plate and the fluorescence measurement was recorded on an excitation of 480 and emission of 521 nm. For further analysis the PE-TB treated aggregates and untreated control aggregates were loaded on 10% SDS-PAGE. The SDS-PAGE was stained with 1% coomassie brilliant blue solution for 10 minutes, after destaining the gel was analysed for presence of higher order aggregated in treated and untreated samples.

### Statistical Analysis

The statistical data for the fluorescence measurement or viability assay was plotted by using either duplicate or triplicate reading. Untransformed (raw) data were analyzed and plotted by SigmaPlot software. The data was analyzed for the significance by unpaired student’s t-test.

## Results

### TB inhibit the Tau aggregation *in-vitro*

The four repeat region in Tau is considered as aggregation prone, VQIINK and VQIVYK hexapeptides present in second and third repeat majorly contribute to the aggregation propensity of Tau. The repeat Tau have basic charge with an isoelectric pH of 9.6 (Fig. 1A). TB is a basic dye belonging to phenothiazine group of compounds (Fig. 1B). The aggregation inhibition property of TB was studied against heparin-induced Tau aggregation. The fluorescence studies suggested that TB efficiently inhibited Tau aggregation in concentration-dependent manner. The inhibition was observed at a concentration of 2 μM and the rate of inhibition increased proportionally with the concentration of TB (Fig. 1C). 40 μM of TB found to inhibit 80% of aggregation (p ≥0.001), suggesting that TB to be a potent molecule against Tau aggregation (Fig. 1D). The electron micrograph of TB treated Tau showed broken and fragile fragment unlikely of untreated sample, comprising of long and thick filamentous Tau aggregates (Fig. 1E-F).

**Figure 1.**
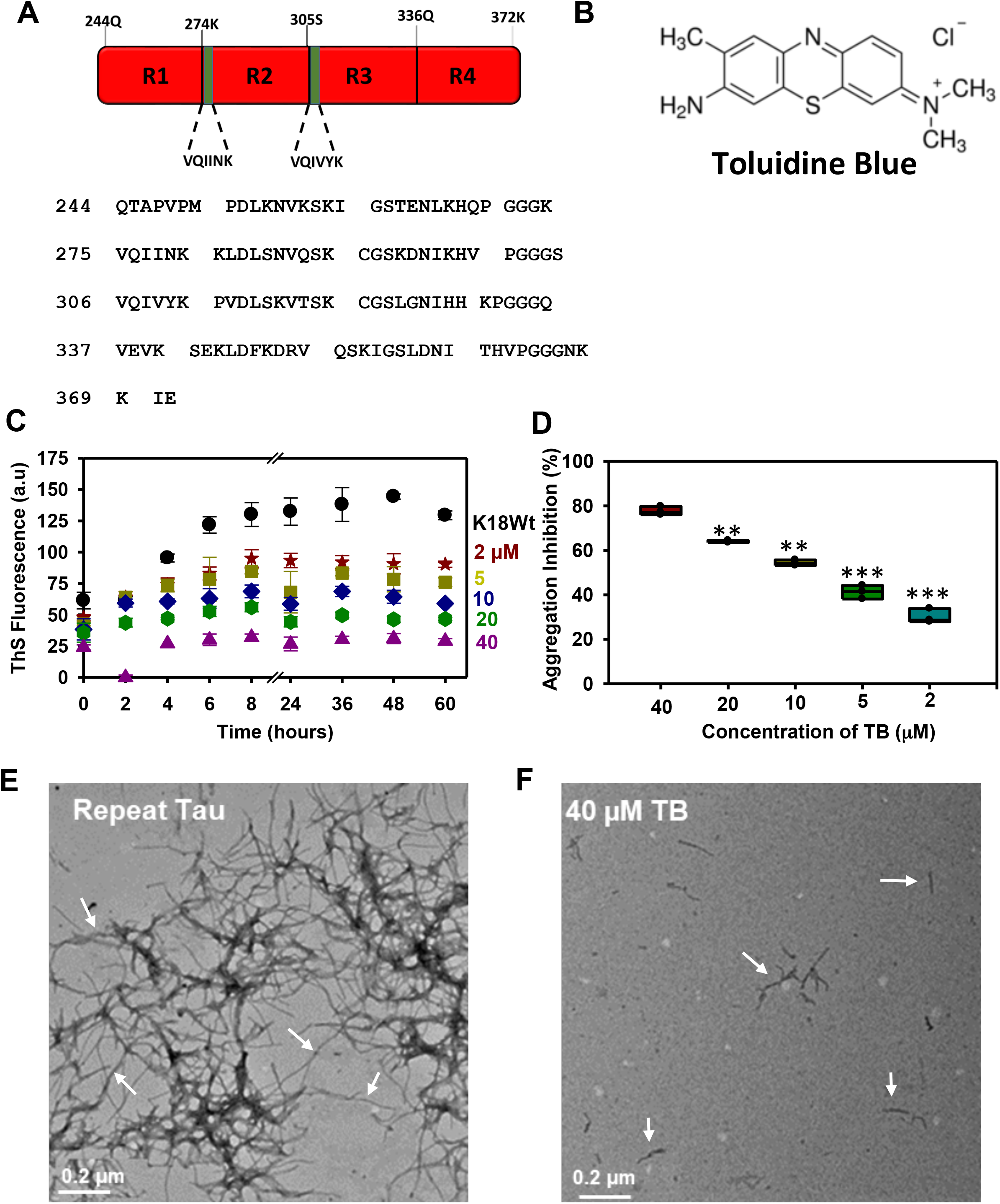
The aggregation propensity of repeat Tau with Toluidine Blue. A) The four repetitive region in Tau are prone for aggregation. The TB is a phenothiazine dye having a basic charge. B Aggregation of repeat Tau in presence of TB was probed by ThS fluorescence. Result suggested the aggregation inhibition of Tau in concentration-dependent manner with respect to TB. TB found to inhibit the aggregation at concentration as low as 5 μM. C) The end point fluorescence analysis to quantify the rate of aggregation inhibition by TB in concentration-dependent manner. E, F) The electron microscopic analysis of aggregated Tau showed long thick filaments, while after treatment the broken filaments were observed. The significance was calculated using unpaired Student t-test in SigmaPlot software. *p<0.05, **p<0.001, ***p<0.0001, the statistical difference between control and treated groups.

### Photo-excited TB dissolved the matured Tau filaments

TB has an absorption maxima of 630 nm, the irradiation of TB at 630±20 nm leads to conversion of TB in photo-excitation form, which generates singlet oxygen species (Fig. 2A). In our study, we exposed TB under a red LED light source for 180 minutes for photo-excitation with irradiance of 9.9*10^6^ Watt/m^2^. Additionally one light control (LC) was also considered to study weather red light alone could have potency to dissolves the Tau fibrils, The Tau treated with PE-TB showed no higher ordered aggregates, which was evidenced by SDS-PAGE (Fig. 2B). Furthermore, ThS assay was carried out for evaluating the disaggregation potency of PE-TB. These results suggested that in comparison to untreated aggregates fluorescence intensity was decreased in PE-TB treated aggregates samples. Moreover red light alone was not effective on Tau aggregates as the LC had similar fluorescence as untreated control (Fig. 2C). The electron microscopy images support the observation that PE-TB efficiently disaggregated the mature repeat Tau fibrils (Fig. 2D-E). Thus, the fluorescence assay data supported the fact that PE-TB disaggregates the pre-mature repeat Tau filament in concentration dependent manner. In our experiment negligible heat changes were observed after 180 minutes of irradiation. These *in-vitro* experiments clearly indicated that PE-TB to be an effective molecule against repeat Tau aggregates.

**Figure 2.**
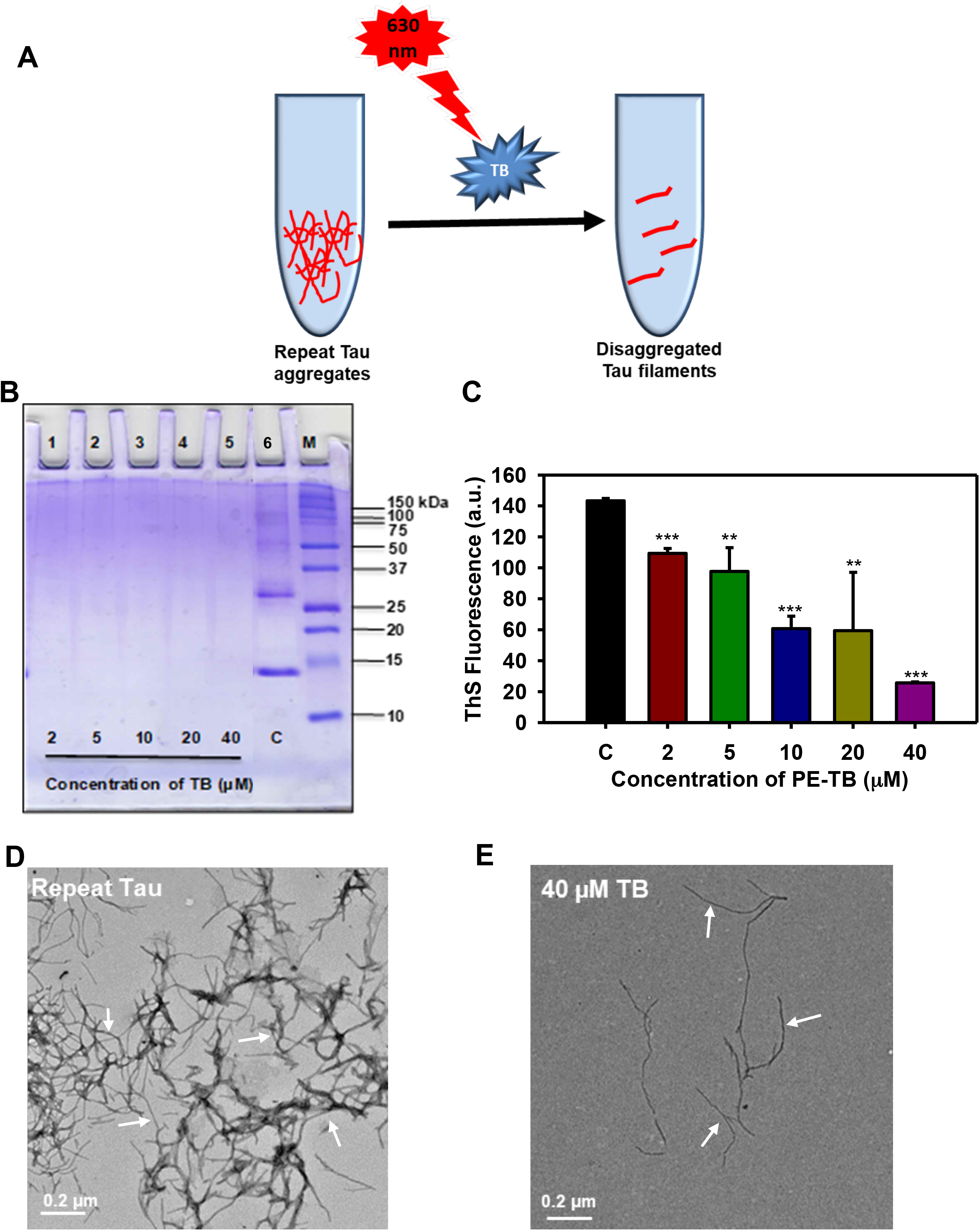
Effect of PDT on repeat Tau aggregates at various concentrations of photosensitizer. A) The repeat Tau aggregates were incubated with TB, which was further irradiated with 630 nm of red light. B) PE-TB (2-40 μM) on repeat Tau aggregates. The results observed from the experiment suggested that PE-TB completely dissolved the higher molecular weight aggregates of repeat Tau. C, D) The treatment with PE-TB generated morphological changes in Tau fibrils. The aggregated Tau has long thick filaments while after the treatment predominantly fragile and broken filaments were observed under electron microscope. E) The ThS analysis showing the disaggregation potency of PE-TB. PE-TB treated aggregates have low fluorescence, which indicated the absence of higher order aggregates in sample. The significance was calculated using Student t-test in SigmaPlot software. *p<0.05, **p<0.001, ***p<0.0001, the statistical difference between control and treated groups.

### The cytotoxicity of higher concentration of TB

The current experiment aimed to study the toxicity for TB and PE-TB against mouse neuroblastoma cells, Neuro2a. The cells were treated with varying concentrations of TB (1-100 μM) for 24 hours. Additionally, TB incubated cells were irradiated with 630±20 nm red light for 10 min with an irradiance of 5.5*10^5^ Watt/m^2^. These results suggested that TB-induced low levels of toxicity than PE-TB. TB found to have minimal toxicity at a concertation of 40 μM where as a concentration of 120 μM induced-toxicity. However, PE-TB was found to be toxic at a concentration of 20 μM (Fig. 3A-B). We speculate that the singlet oxygen produced by PE-TB might lead to generation of toxicity. Furthermore at concentration of 80 μM and above TB was observed to be internalized in neuronal cells (Fig. 3C). Interestingly, we observed that at a concentration of 80 μM and above PE-TB was internalized in neuronal cells. Here we speculate that accumulation of internalized PE-TB could also be lead to toxicity. The overall results suggested that TB has low toxicity at even at higher concentrations.

**Figure 3.**
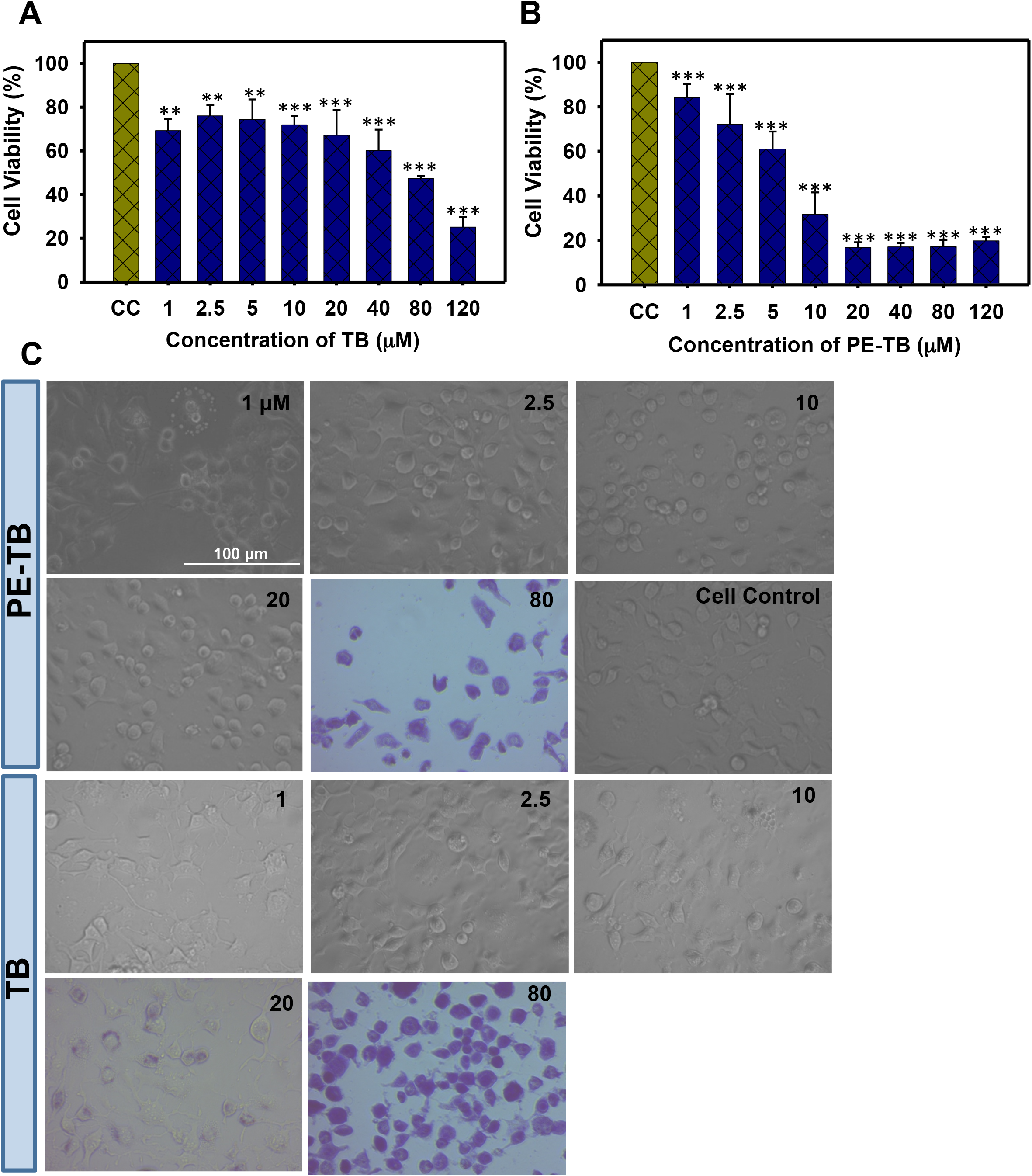
The effect of higher concentration of TB was observed on N2a cell viability. A) Photo-excited TB was found to be toxic at higher concentrations. B) The effect of non-photo-excited TB was observed on N2a cells. The data suggested that TB was toxic at higher concentrations. C) Morphological analysis of TB treated cells. 80 μM TB was found to be internalized in cells. The significance was calculated using student t-test in SigmaPlot software. *p<0.05, **p<0.001, ***p<0.0001, the statistical difference between control and treated groups.

### Modulation of cytoskeleton by PE-TB

Tubulin is basic unit of cytoskeleton, which polymerizes leading to formation of microtubules. The effect of PE-TB on cytoskeleton was observed by immunofluorescence studies. Neuro2a cells were incubated for 24 hours with varying concentrations of PE-TB (0.5 and 50 μM). The immunofluorescence studies suggested that as compared to untreated cell control, cell treated with 0.5 μM PE-TB have high fluorescence intensity of tubulin, whereas no differential results were observed in cells treated with 50 μM of PE-TB treatment (Fig. 4A-B). The quantification of fluorescence images suggested minimal increase in tubulin intensity. Tubulin is the basic unit of microtubules thus the modulation of tubulin intensity suggested modification of cytoskeleton after PE-TB treatment (Fig. 4C). Although the morphological changes in cell after PE-TB treatment was not appreciably significant as minimal changes in elongated dendritic extensions were observed in treated cells (Fig. 4D). Moreover, to analyse the effect of TB and PE-TB on cytoskeleton we analysed the actin modulation (Fig. 5A). Our studies indicated that TB treatment induces actin structure in cells, as the cells were observed to have actin-rich “Lamellipodia” structures after TB exposure. We also observed the elevated numbers of fine hair-like protrusions or “Filopodia” in cells. While, the cells treated with PE-TB observed to have lamellipodia structures, on contrary filopodia structures were not apparent after PE-TB treatment (Fig. 5B). Furthermore, we studied the End-binding protein-1 (EB-1) expression in neurons. These result suggested that TB and PE-TB treatment enhance the levels of EB1, which suggest that TB and PE-TB treatment might accelerate the microtubule polymerisation in cells (Fig. 6A-B). The immunoblot analysis was carried out for the TB and PE-TB treated samples. The results of immunoblot suggested that 0.5 μM of TB and PE-TB treatment elevated the EB1 levels in cells. The elevated levels of EB1 indicated that there could be an increased rate of microtubule polymerization after the TB and PE-TB treatment (Fig. 6C).

**Figure 4.**
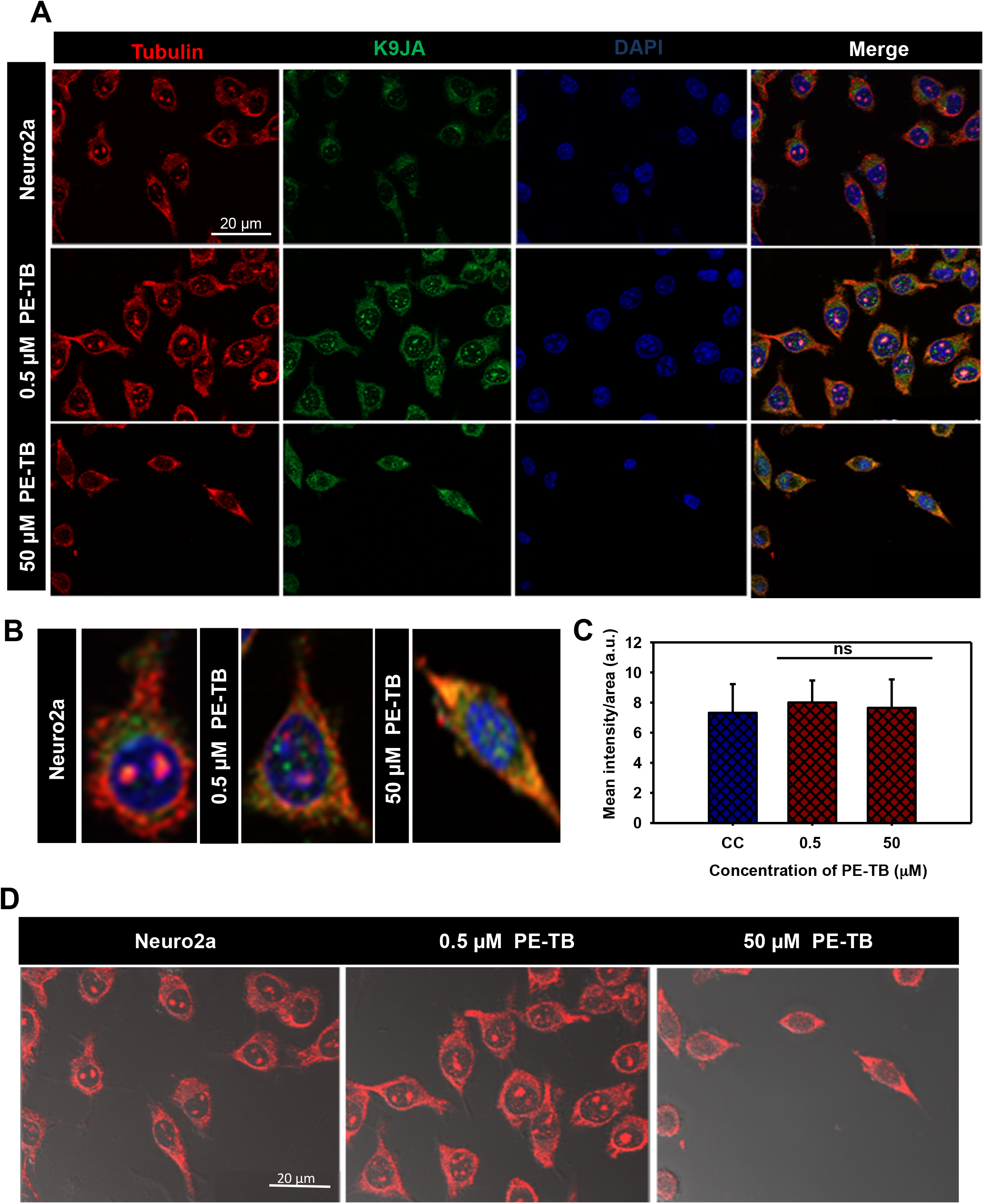
The effect of PE-TB on cytoskeleton. A) The effect of PE-TB on cytoskeleton was studied by immunofluorescence. The cell treated with various contrition of PE-TB (0.5 μM and 50 μM) showed modulation of tubulin intensity. Increase in tubulin intensity was observed after treatment of 0.5 μM PE-TB treatment. B) The single cell images showing the differential level of tubulin in PE-TB treated cells. C) The quantification of immunofluorescence images suggested that intensity of tubulin increased after PE-TB treatment to cells. D) Images including DIC showing the morphological changes after PE-TB treatment of Neuro2a cells.

**Figure 5.**
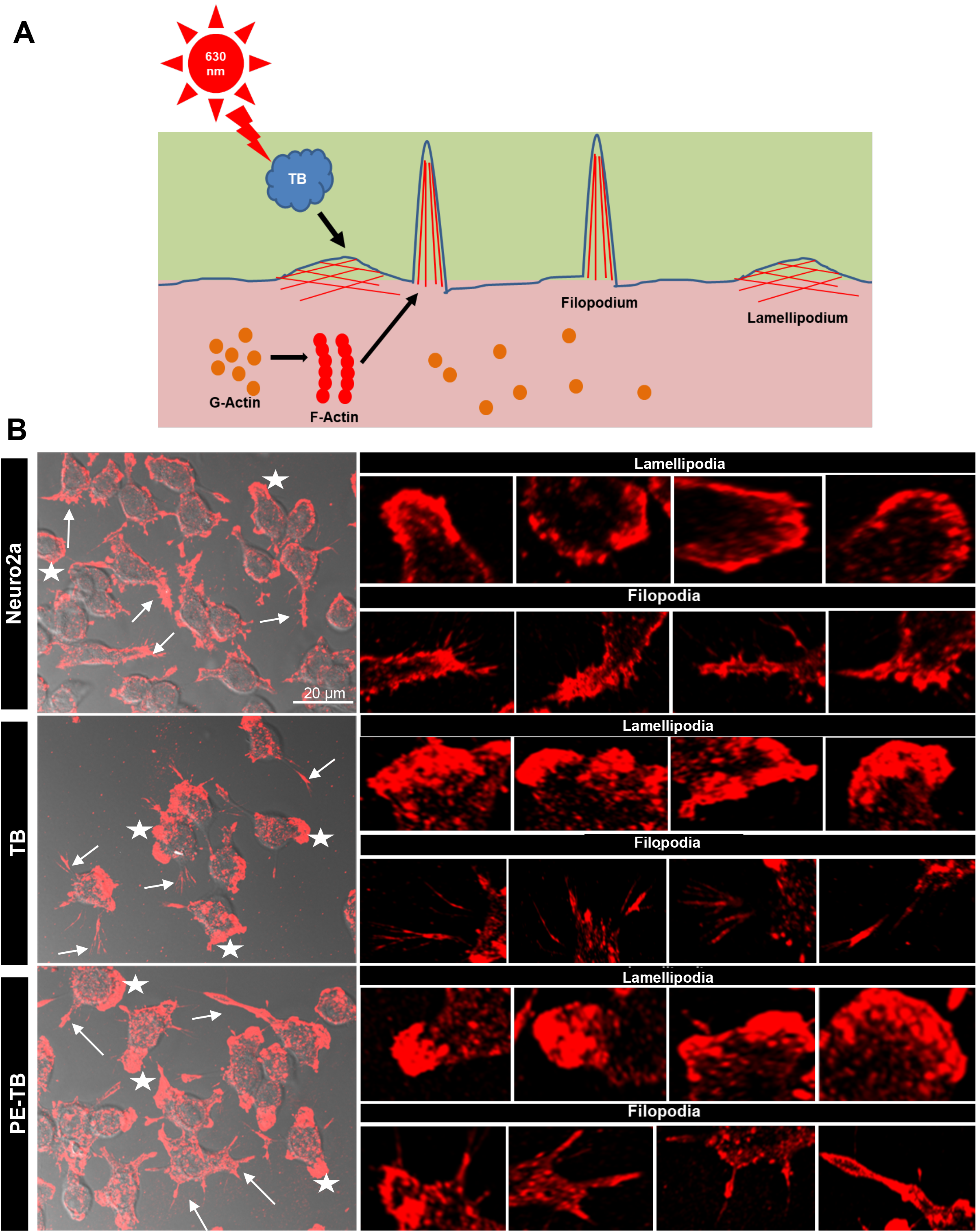
TB induces modulation in cytoskeleton. A) Schematic diagram for the alteration of actin cytoskeleton after TB and PE-TB treatment. B) The immunofluorescence images showing the changes in actin cytoskeleton, the filopodia (arrow marked) and lamellipodia (star marked) structures were modulated after TB and PE-TB treatment.

**Figure 6.**
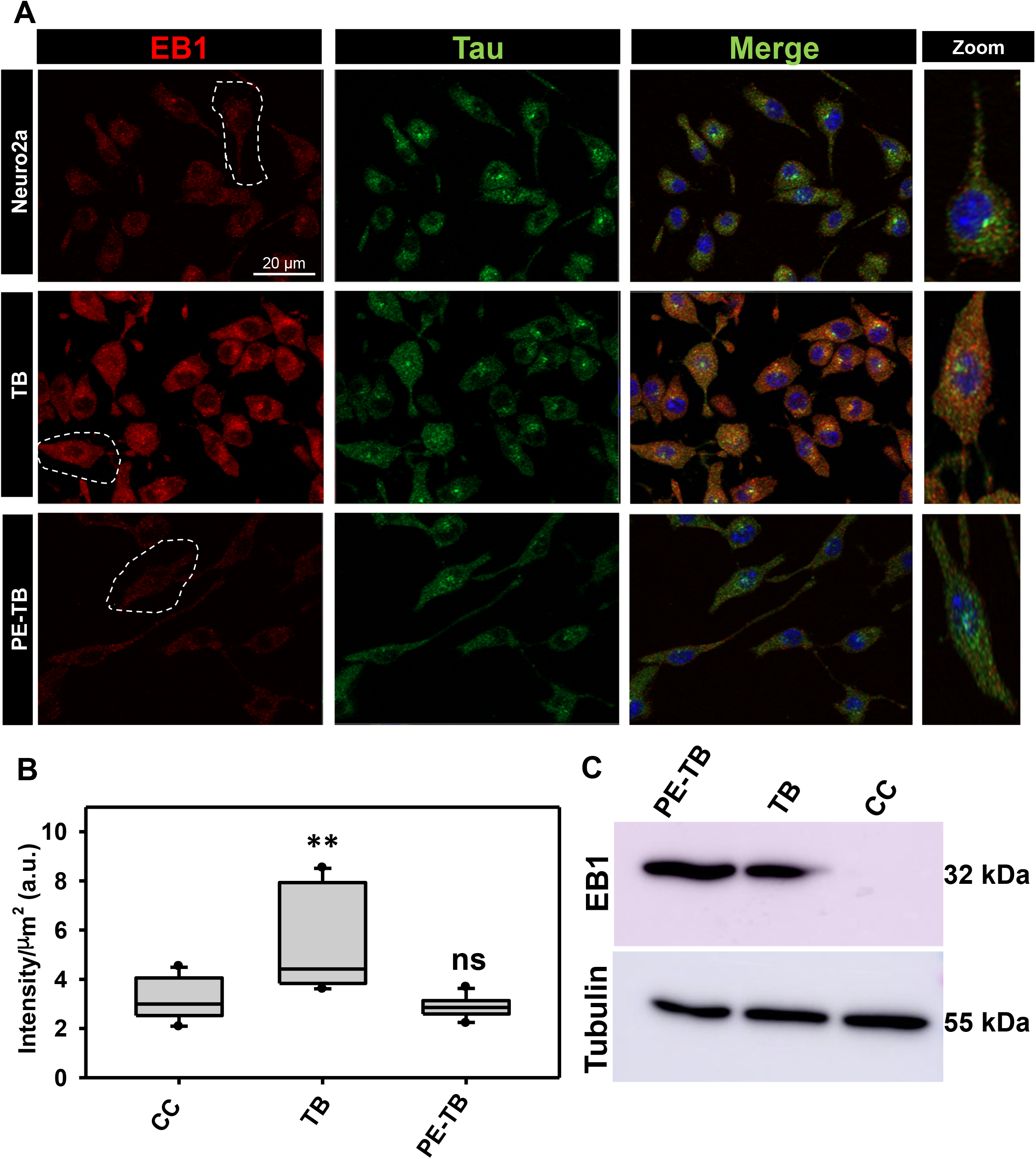
TB and PE-TB induces changes in EB1 expression. A) The immunofluorescence images showing the changes in EB 1 levels after TB treatment. B) The quantification of immunofluorescence images suggested the TB increases the EB1 levels in cells. The quantification was done by using Zen 2.0 blue software. C) The western blot analysis of the studying the expression levels of EB1 in cells. The western blot suggested that TB and PE-TB modulated the EB 1 levels in cells as compared to control.

## Discussion

The pathological state of Tau leads to formation of Tau aggregates, several factors including post-translational modification, reactive oxygen species and mutations result in generation of Tauopathies(Iqbal *et al.* 2005; Iqbal *et al.* 2016). Several studies have been done for screening of the compounds against Tauopathy (Gorantla *et al.* 2018). Various classes of dyes were tested for their potency against neurodegeneration. The aggregation inhibition potency of methylene blue for Tau has been already reported illustratively. Xanthene dye such as erythrosine B was reported to reduce the Amyloid-β-mediated toxicity in cells. Photo-excited Rose Bengal was found potent in inhibiting the Amyloid-β aggregation in liver cells. Furthermore the sulfonated dye Congo red was reported to attenuate the Amyloid-β aggregation. Phenothiazine classes of dye are reported to have therapeutic potency against numerous diseases (Marshall *et al.* 2019). Methylene blue and its derivatives have been reported to be neuroprotective molecules (Oz *et al.* 2009; Poteet *et al.* 2012). TB is a known basic dye used in histological staining but the medicinal properties of TB has not been studied illustratively (Sridharan & Shankar 2012). Here we have studied the efficiency of TB against Tau aggregation and our observations suggested that as its parent compound methylene blue, TB was also efficient in inhibiting the aggregation of repeat Tau. Recently researches have investigated that irradiation plays a crucial role in treatment of AD. The intranasal red light probe, which irradiate red light on brain areas found to show significant recovery in mouse model of AD (Dimauro *et al.* 2008). Similarly, light-based intracranial implants have been patented for treatment of Parkinson’s disease patients (Dimauro *et al.* 2007). Moreover, EGCG irradiated with near infra-red light found to reduce the Amyloid-β plaques deposition in neuronal cells (Johnstone *et al.* 2016). TB is a well reported photosensitizer, which was found efficient as an antibacterial molecule (Sharma *et al.* 2008). In our studies we irradiated TB with 630 nm red light to study its effect on Tau aggregates. TB also reduced the Amyloid-β aggregation but the effect of but TB was found ineffective against Tau hyperphosphorylation (Yuksel *et al.* 2018). We found that PE-TB potentially disaggregated the pre-formed Tau filaments. The PE-TB led to generation of singlet oxygen species, thus we speculate that singlet oxygen contributes in disaggregation potency of TB. The SDS-PAGE and electron microscopic studies clearly indicated the disaggregation of Tau filaments by PE-TB. ThS dye binds specifically to protein aggregates thus by tracing the ThS fluorescence, we could observe the extent of protein aggregation. In our results the PE-TB treated aggregates showed low ThS fluorescence as compared to untreated aggregates, which indicated that PE-TB disaggregated the repeat Tau filaments. *In-vitro* studies including SDS-PAGE, electron microscopy and ThS fluorescence clearly advocated that PE-TB treatment dissolved the mature Tau filaments. The phenothiazine dye-induced low levels of toxicity in cells, but on photo irradiation the dyes induced toxicity in cells due to generation of singlet oxygen species (Wainwright *et al.* 1997). In our work, we found that TB was not significantly toxic to cell even at a high concentration of 40 μM but the PE-TB reduced the viability of cell at a concertation of 20 μM. These results suggested that TB at lower concentration could be considered as biocompatible. The potency of photo-irradiation in modulating the cytoskeleton has been reported in human hepatoma cells (Liu *et al.* 2006). The reports suggested that irradiation inhibited the cell growth by modulating the cytoskeleton. In our results, we observed that PE-TB treatment minimally increases the tubulin intensity in neuronal cells, which indicated that PE-TB could modulate cytoskeleton structure. The actin cytoskeleton plays crucial role in cell motility and cell synapse formation(Luo 2002). The lamellipodia are the actin-rich structures and the modulation in lamellipodia led to change in cell motility and adhesion(Small *et al.* 2002). Similarly the modulation in filopodia structure alter the cell synapse formation and motility(Mallavarapu & Mitchison 1999). Thus, the alterations in actin network after TB and PE-TB treatment led us to conclusion that the dye has potency to modulate cell cytoskeleton networks. Hence, the overall results suggest that PE-TB could have a therapeutic potency against Tauopathy (Fig. 7).

**Figure 7.**
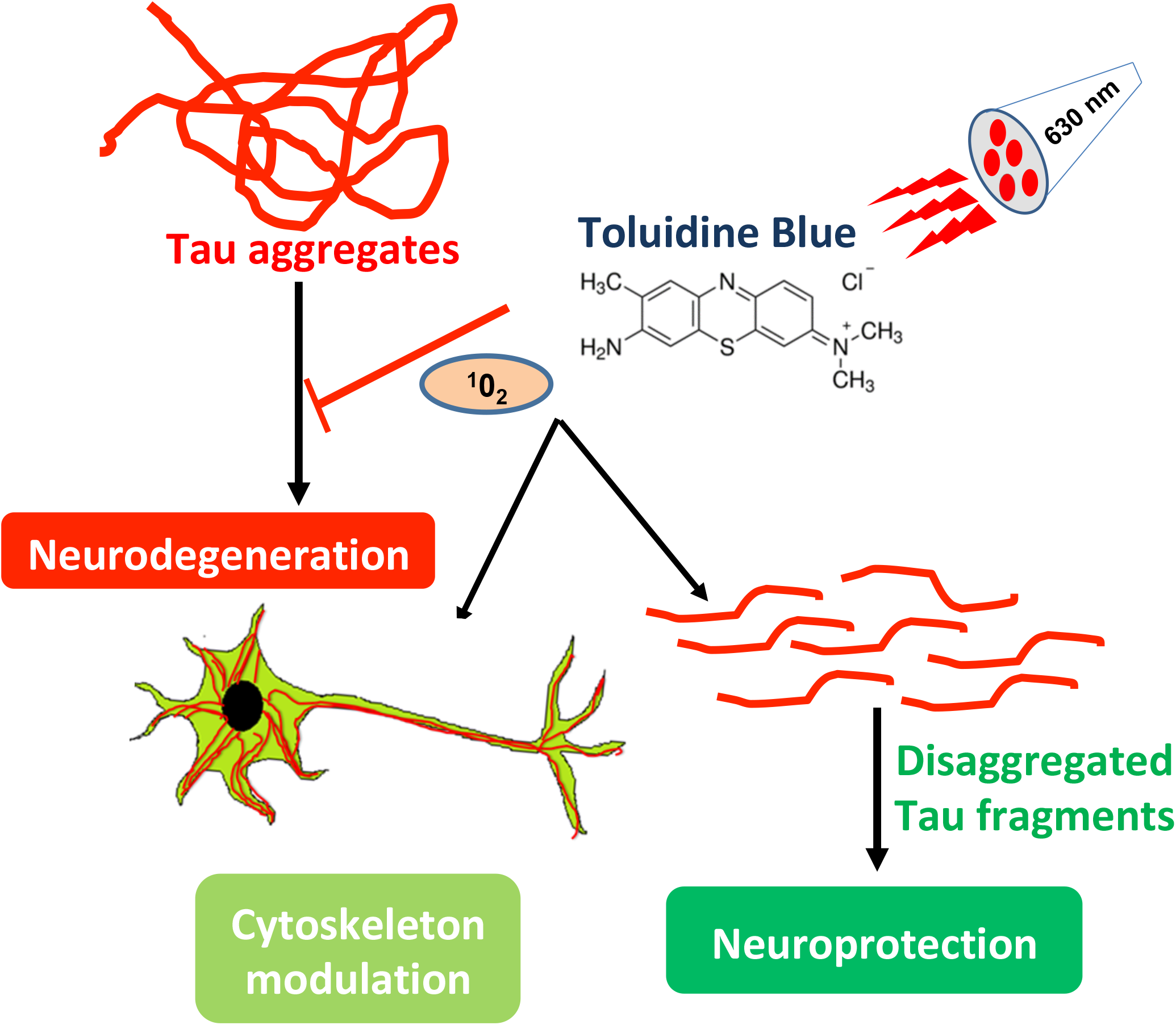
The TB and PE-TB dissolve the Tau fibrils. Tau aggregates are one of the hallmarks of AD. Accumulation of Tau aggregates lead to generation of neurodegenerative diseases. TB on irradiation with 630 nm red light gets converted to PE-TB. PE-TB disaggregates the pre-formed Tau and modulate the cytoskeleton. Thus, PE-TB could be a potent molecule against Tauopathy.

## Conclusions

Tau aggregation leads to generation of toxic paired helical filaments, which is hallmark of AD. The molecules having potency to inhibit the Tau aggregation are being considered as a therapeutic molecules in AD. In our study TB and PE-TB were observed to be potent against Tau aggregation. We observed a distinctive modulation of actin, tubulin networks and EB1 levels after TB and PE-TB treatment, which clearly suggested that TB and PE-TB have potency against the cytoskeleton deformities. Thus, from the present work we are stating that TB could be a potent molecule against Tauopathy.

## Acknowledgements

The authors acknowledge CSIR-National Chemical Laboratory for the instrumentation central facility. This project is supported by grant from the in-house, National Chemical Laboratory-Council of Scientific Industrial Research-(CSIR-NCL) MLP029526. Tushar Dubey and Nalini Vijay Gorantla acknowledges the fellowship from University of Grant Commission (UGC), India.

## Author Information

Tushar Dubey, Nalini Vijay Gorantla and Subashchandrabose Chinnathambi

**Neurobiology Group, Division of Biochemical Sciences, CSIR-National Chemical Laboratory, Dr. Homi Bhabha Road, 411008 Pune, India**

Tushar Dubey, Nalini Vijay Gorantla and Subashchandrabose Chinnathambi

**Academy of Scientific and Innovative Research (AcSIR), 110025 New Delhi, India**

## Author Contributions

S.C designed the experiments. T.D and N.V carried out the experiments. T.D and S.C analyzed the data. S.C. conceived the idea of the project, provided resources, supervised and wrote the manuscript. All authors contributed to the discussions and manuscript review.

## Corresponding Author

Correspondence and requests for materials should be addressed to **Prof. Subashchandrabose Chinnathambi. Email:**

**Telephone: +91-20-25902232, Fax. +91-20-25902648.**

## Competing Financial Interests

The authors declare no competing financial interest.

## Abbreviations

AD, Alzheimer's disease; PHFs, Paired Helical Filaments; NFTs, Neurofibrillary Tangles; TB, Toluidine Blue; PE-TB, Photo-excited TB; MB, Methylene Blue; PDT, Photodynamic Therapy; ThS, Thioflavin S; MTT, Methylthiazolyldiphenyl-tetrazolium bromide; SEC, Size Exclusion Chromatography; BCA, Bicinchoninic acid; DMSO, Dimethyl sulfoxide, SDS-PAGE, Sodium dodecyl sulfate-polyacrylamide gel electrophoresis; TEM, Transmission electron microscopy; BSA, Bovine serum albumin.

